# cT-DNA in *Linaria vulgaris* L. is multicopy, inverted and homogenized

**DOI:** 10.1101/615328

**Authors:** Ivan A. Vladimirov, Olga A. Pavlova, Dmitrii E. Polev, Denis I. Bogomaz

**Affiliations:** Beagle Ltd, 152-1-77, Bukharestskaya str., Saint-Petersburg, Russia, http://biobeagle.com/; Peter the Great St.Petersburg Polytechnic University, 29, Polytechnicheskaya str., Saint-Petersburg, Russia, http://english.spbstu.ru/; Saint-Petersburg State University, 7/9, Universutetskaya emb., Saint-Petersburg, Russia, https://spbu.ru/

**Keywords:** cT-DNA structure, *Linaria vulgaris*, inverted repeats, microsatellite markers, concerted evolution

## Abstract

The phenomenon of evolutionary fixation of agrobacterial sequences (cT-DNA or cellular transferred DNA) in plant genomes is well known in nature. It was previously considered, that all of cT-DNA-containing species, except *Linaria vulgaris*, have multiple inverted cT-DNA repeats. Deep studying of general features of cT-DNA brings us closer to understanding the causes and mechanisms of its fixation in plants genomes. We combined multiple long-range PCR with genome walking for studying extended structure of cT-DNA. Using digital PCR method, we estimated copy number of cT-DNA elements. NGS with low covering allows us to develop a set of microsatellite markers, also used for copy number estimation. According to new data, cT-DNA elements in *L. vulgaris* form an inverted complex repeat of two simple direct repeats. After cT-DNA integration, cT-DNA sequence duplication events took place at least two times. The phenomenon of concerted evolution of cT-DNA sequences as well as some details of this process have been shown for the first time.

We have shown, that *L. vulgaris,* as well as other cT-DNA containing species, has inverted structure of repeats. This fact indicates possible existence of some general causes and mechanisms of cT-DNA fixation in plant genomes during evolution.

## Introduction

*Linaria vulgaris* is an example of a plant containing a vertically inherited sequence of *Agrobacterium rhizogenes* T-DNA in the genome (cT-DNA or cellular T-DNA). It is a rare phenomenon, currently known only in *Linaria* (Matveeva et al. 2012; Pavlova et al. 2014), *Ipomaea* (Kyndt et al. 2015) and *Nicotiana* genera (White et al. 1983; Intrieri and Buiatti 2001). The fixation of cT-DNA sequences in genomes carriesa pathogenic program of genetic colonization (Tzfira and Citovsky 2006). The mere fact of such plant survival is of great interest. The intactness of several cT-DNA genes, as well as the observed expression of some of them, makes it necessary to discuss the function of cT-DNA in the genomes of such plants (Meyer et al. 1995; Frundt et al. 1998; Aoki and Syono 1999, 2000; Suzuki et al. 2002). During the evolution of the genus *Nicotiana* integration of cT-DNAs occured several times in a row (Chen et al. 2014). This fact also serves as an indirect confirmation of a specific function of these cT-DNAs.

*Linaria* plants derived T-DNA from the mikimopine strain of *Agrobacterium rhizogenes*, which is similar to the present “1724” strain (Matveeva et al. 2012; Pavlova et al. 2014). According to the published data, the cT-DNA insert integrated into the transposon and organized a direct repeat of two cT-DNA units (Matveeva et al. 2012; Pavlova et al. 2014).

## Results

This work was started to clarify the *L. vulgaris* cT-DNA structure. Earlier (Matveeva et al. 2012), it was shown that the T-DNA sequence in *L. vulgaris* is an imperfect direct repeat of two T-DNA elements. (We call the “element” a form of a single T-DNA sequence which is found in the agrobacterial Ri-plasmid). However, the sequence EU735069.2 presented in Matveeva et al. 2012 was obtained without taking into account the possibility of high copy number of T-DNA. In this regard, we have conducted a more detailed study of the T-DNA sequence by sequencing Long-Range-PCR products obtained from unique primers. Unique primers were designed for the sites where elements join with each other and with plant DNA. The sequences of sites for elements joining were determined using GenomeWalker (Siebert et al. 1995). Six unique sites for elements joining were sequenced. Three of them (we call them β, β’, γ) are formed by direct repeats, and three (ω, o’, φ) - by inverted repeats (Figure 1) shown for *L. vulgaris* for the first time. Theprimers selected for joining regions produced single target PCR-products, so sequences of joining regions could not be PCR-artifacts. The relative positioning of the elements was established using the Long-Range-PCR. The resulting PCR products containing sequences of individual elements were sequenced (Table 3).

**Table 1.**
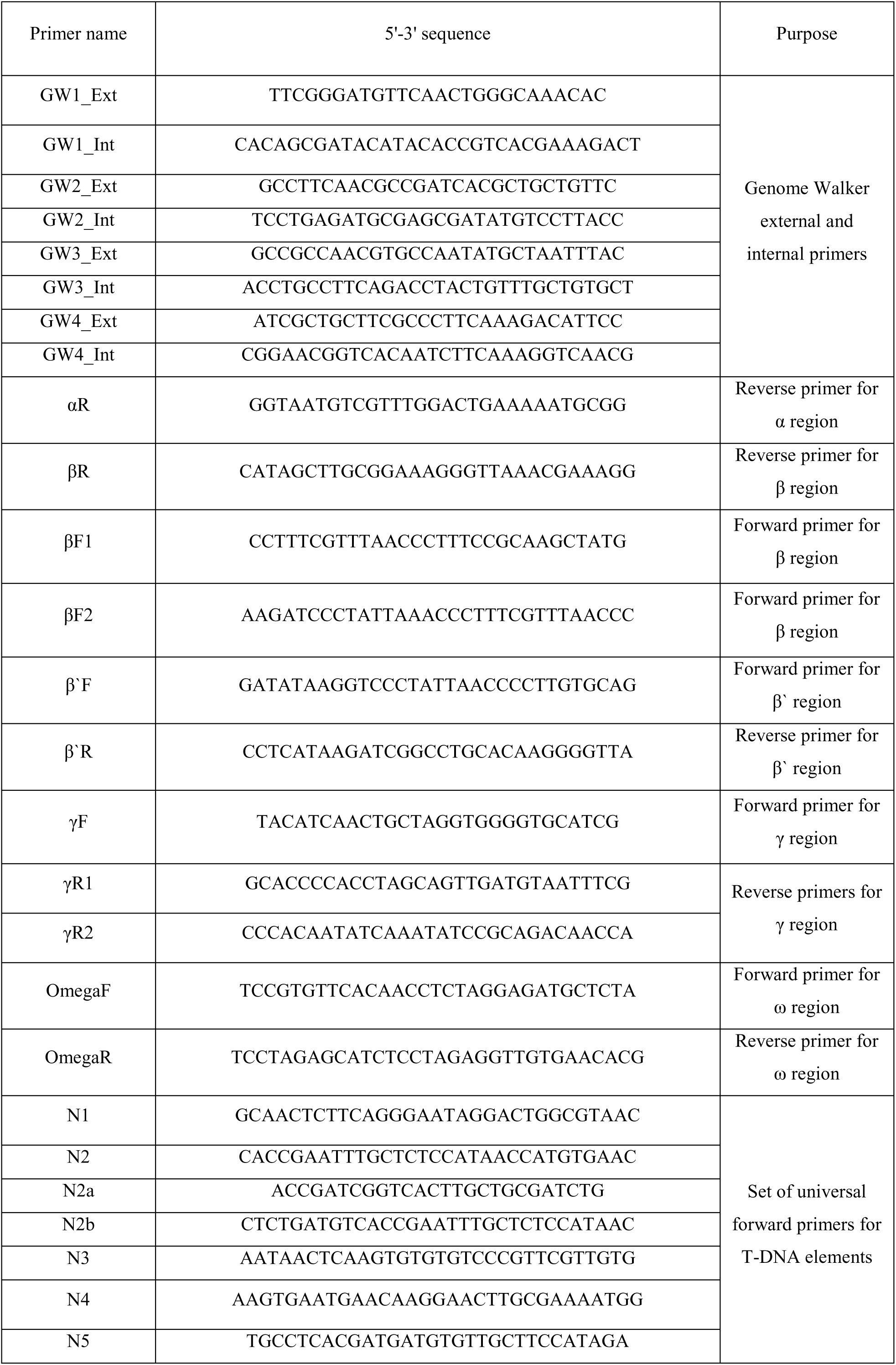
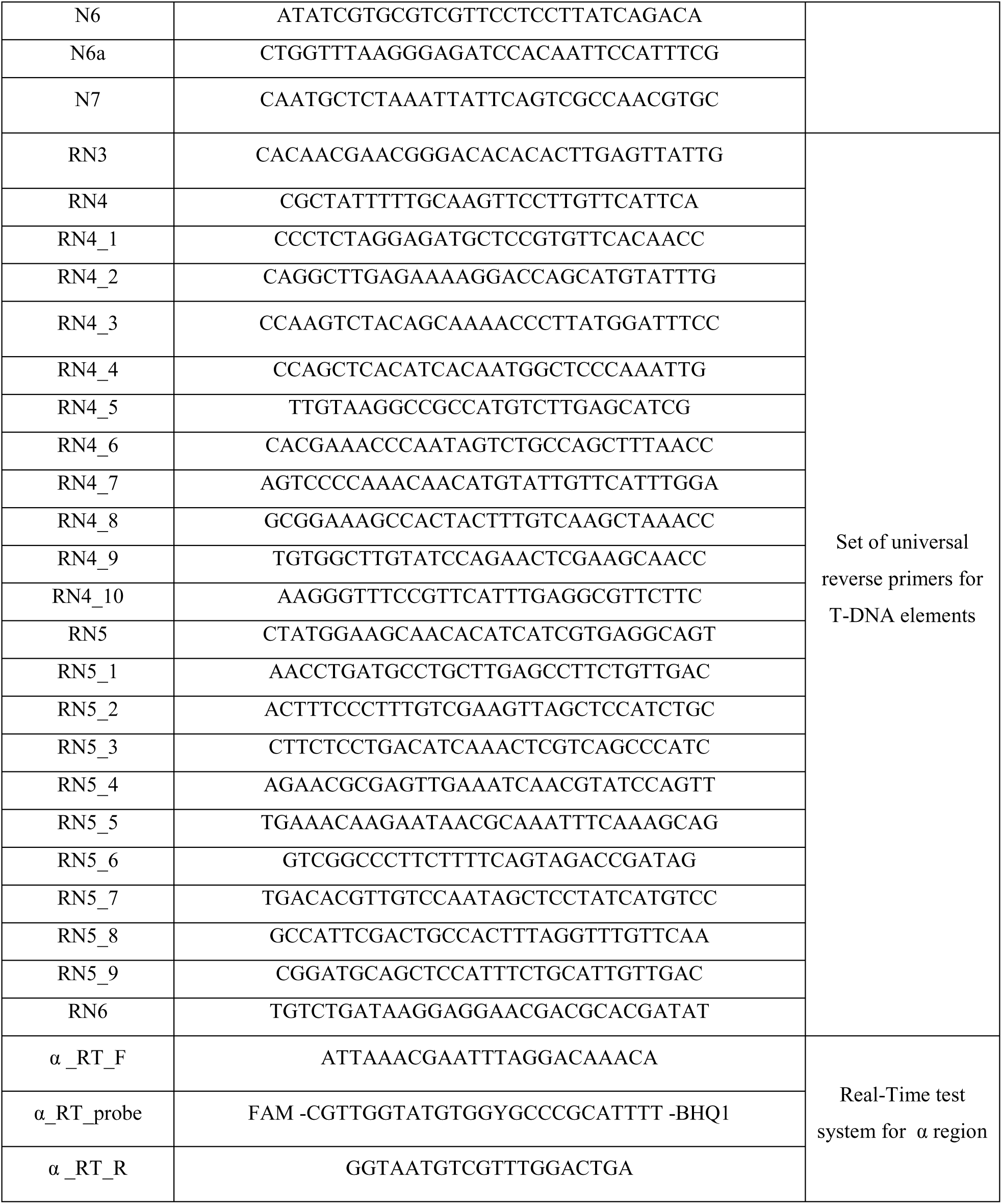
Primers list.

**Table 2.**
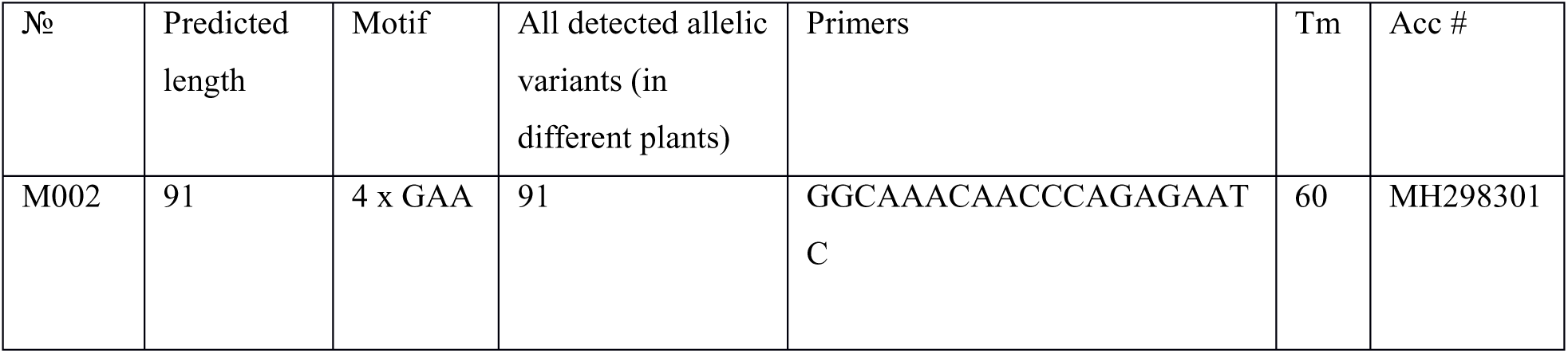
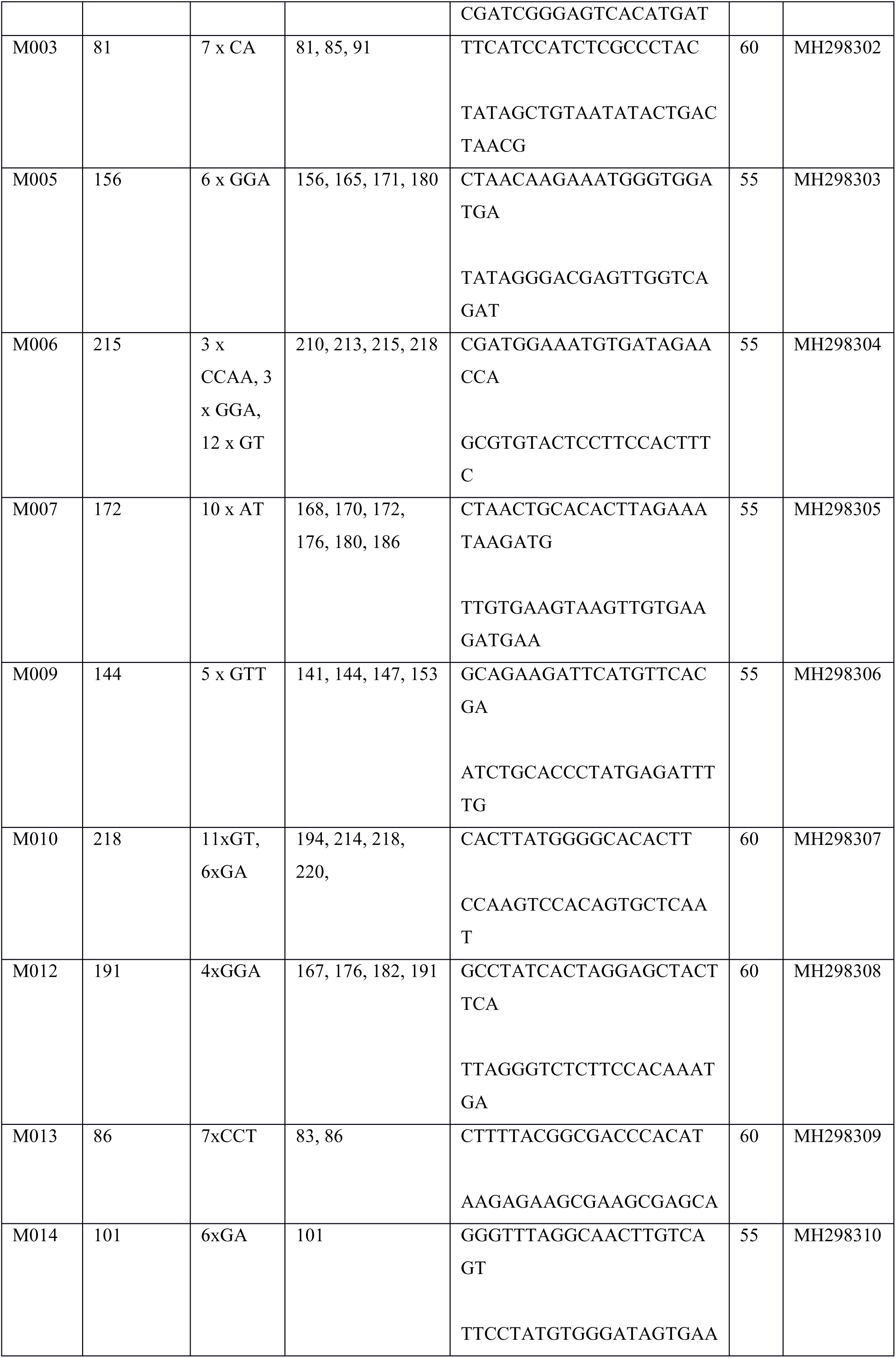
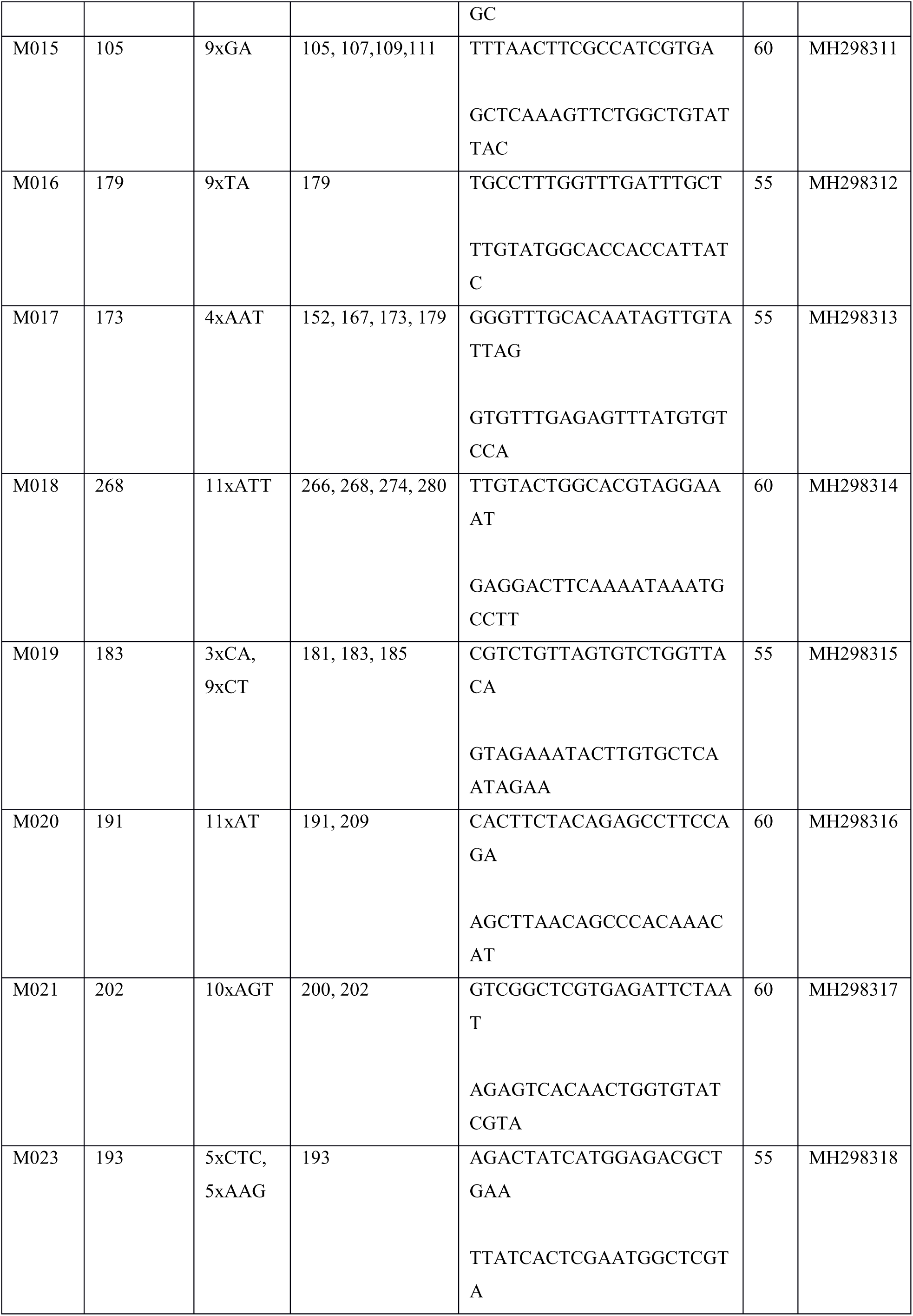
Microsatellites markers list

**Table 3.** Accession numbers.

| Sequence name | Acc. number in NCBI |
| --- | --- |
| $\gamma$ sequence | KR612321.1 |
| 9.13 | MH107074 |
| 18.2 | MH107075 |
| 20.1 | MH119454 |
| 20.2 | MH119451 |
| 20.3 | MH119450 |
| 20.4 | MH119453 |
| 20.5 | MH119452 |
| 89.1 | MH119458 |
| 94.7 | MH119455 |
| 106.10 | MH119459 |
| 135.11 | MH119456 |
| 160.1 | MH119460 |
| 206.2 | MH119457 |
| right border | MH298319 |

**Figure 1.**
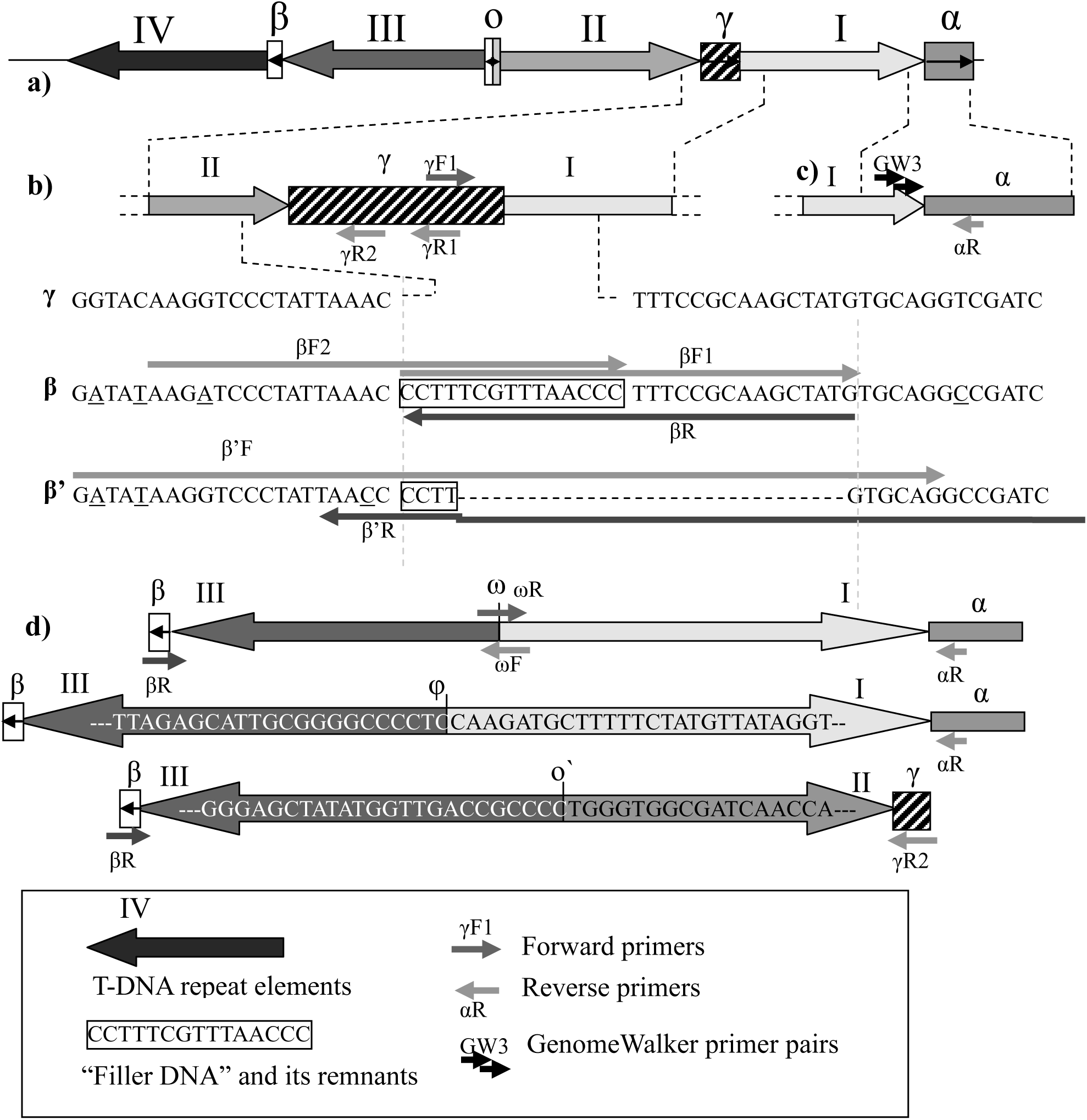
The construction of sites for elements joining and the positions of primers: a) Reconstructed primary structure of the T-DNA block in *L.vulgaris*; b) differences in the sequence between the sites γ, β, β′; c) structure of α (junction of element I with plant DNA); d) The structure of joining sites formed by inverted repeats (ω, φ, o′).

Full-size blocks of fragments were sequenced (Fig.2b), as well as individual fragments (20.2, 160.1, 20.5, 18.2, 20.1, 135.11, 206.2) which could be their allelic variants.

Analysis of the sequences gave us an opportunity to investigate T-DNA structure in *L. vulgaris* (Fig. 2a). The block of T-DNA elements is represented by a complex inverted repeat consisting of two simple direct repeats, with variants of deletions of various sizes. The positions of the elements are marked with Roman numerals (I-IV from right to left), and the sites of joining between them with Greek letters. The most intact of all joining sites is γ between elements I and II (Fig. 1b). β sequence is located between III and IV elements and it differs from γ by a large deletion. Instead of the deleted fragment, there is a sequence of 15 bp long, which is not present in γ and looks like a typical “filler DNA” (Neve et al. 1997), which formed directly upon the transformation during integration of T-DNA into the genome. The β sequence has a short allelic variant with nested deletion (β’) which also contains traces of “filler DNA” (Fig. 1b). Docking sites formed by inverted repeats (ω, φ, o’) do not contain filler DNA. These sites seem to be a result of big deletions which occurred after transformation act (Fig. 1d).

**Figure 2.**
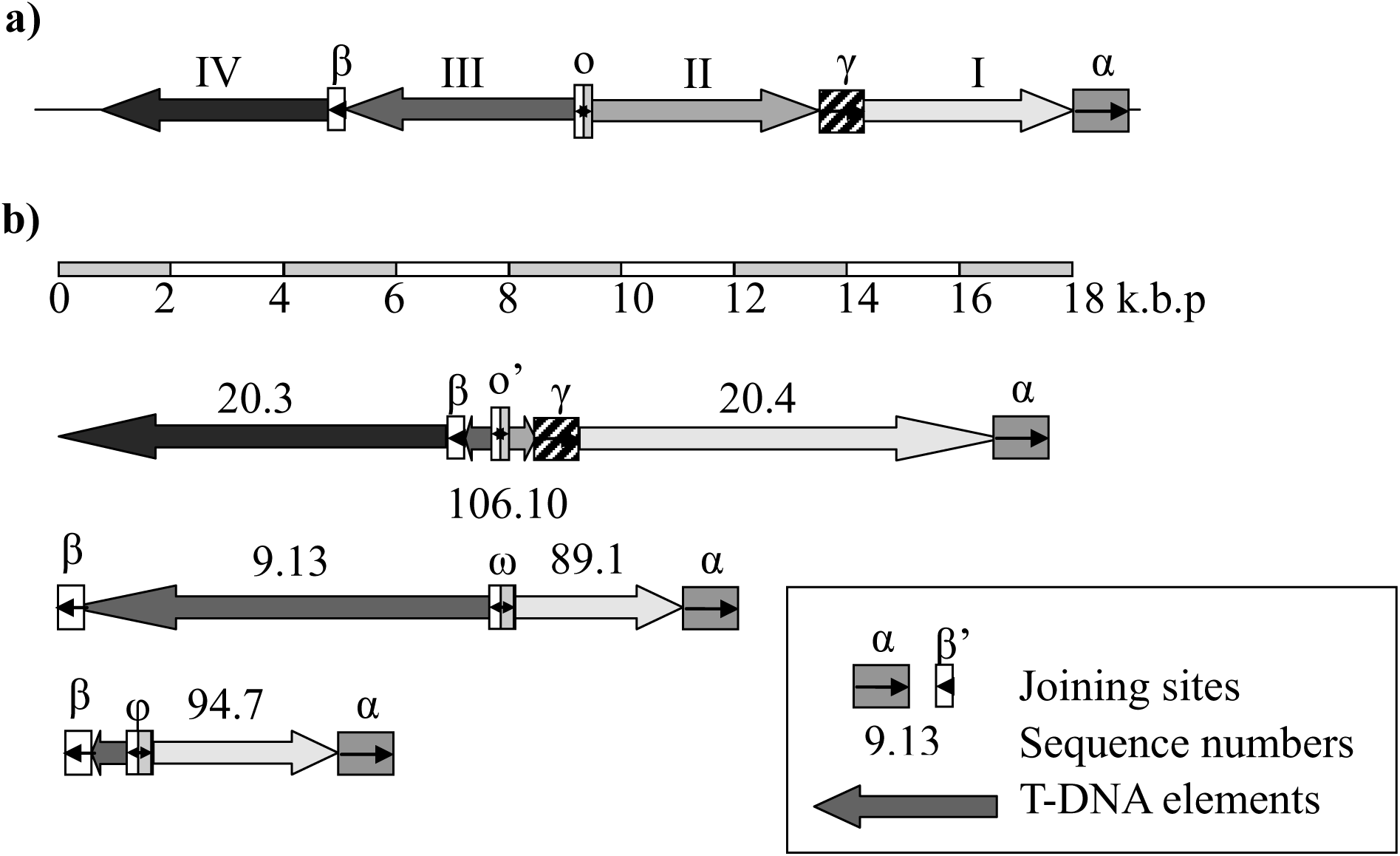
a) Reconstructed primary structure of the T-DNA block; b) Structure of the sequenced T-DNA blocks.

In addition, we found out that:

1) The number of variants of the sequences for almost all positions is more than two (Fig. 2, b). As far as *L.vulgaris* is diploid (Tandon, Bali, 1957), and the number of alleles of a single-copy sequence cannot exceed two, this means that there are several such T-DNA blocks in the *L.vulgaris* genome. In position III five variants of individual sequences corresponding to at least 3 blocks (94.7, 106.10, 20.1, 9.13, in Fig. 2b and 206.2) were detected.

2) Four different variants of the sequences containing the site of joining of T-DNA with plant DNA were found (20.4, 89.1, 94.7- Fig. 2b and 160.1), and the integration site (α) in all four cases was identical.

The analysis of the state of ORF’s from each element was carried out. It was found that all sequenced T-DNA genes except *orf13a* were pseudogenized. Premature stop codons and deletions leading to a reading frame shift were detected in all genes but not in *orf13a*. Gene *orf13a* was found between the *orf13* and *orf14* genes, and it is potentially intact in the sequences 20.1, 9.13 and 206.2.

To clarify cT-DNA copy number digital PCR was carried out. The test system for the α-sequence from cT-DNA was designed. Since *Linaria* genome is not well studied and single copy genes are unknown we decided to use microsatellite markers (previously selected) as reference sequences. Genome sequencing with low coverage was performed using Ion Torrent System for searching candidate sequences for microsatellite markers. Twenty three suitable reads with repeats were selected and primers were designed. All primers were tested on plants DNA from geographically distant regions (Peterhof, Russia and Hakassiya, Russia). Then fragment analysis of PCR products was performed to determine suitability of microsatellite markers. Suitability criteria were: 1) the presence of no more than two peaks of different lengths in each plant studied (indirect confirmation of single copy number); 2) the presence of a at least one peak of the sequence of predicted length in plants sequenced by Ion Torrent System. The list of markers that satisfied all requirements is given in Table 2. The TaqMan probes were designed for markers M006, M009, M010, M014, M015 and M019., Effectiveness of the test systems was estimated by real-time PCR. The M006 marker was excluded since it was multi-copy. The most efficient system based on the M009 marker was selected from the remaining ones. Using a digital PCR the copy number of the α-sequence from T-DNA was compared to the M009 marker. A ratio of 2.19 (1528 copies of the α sequence in 4590 cells and 697 copies in 4590 cells in M009) was obtained.

## Discussion

We have shown an inverted structure of repeats for *L. vulgaris* cT-DNA. Such orientation of the elements during the integration into the genome is often observed (but not necessary). Neve (1997) has shown that the frequency of inverted repeats occurrence depends on the type of T-DNA and varies from 25 to 50% of all cases of multiple elements integration. In addition, there is also a possibility of single T-DNA elements integration. However, the vast majority of known cT-DNAs (TA, TB, TE, TC, TD in *Nicotiana* plants (Chen et al. 2014) and IbT-DNA1 from *Ipomaea batatas* (Kyndt et al. 2015) form exactly inverted repeats. In this work the same organization of cT-DNA (as inverted repeat) in *Linaria vulgaris* genome was first demonstrated.

Given the digital PCR data which demonstrate at least two insertions and taking into account the number of variants of the elements, we can conclude that there are three cT-DNA loci per haploid genome.

Different variants of the sequences containing the same joining site of cT-DNA with plant DNA (20.4, 89.1, 94.7, Fig. 2, b and 160.1 as allelic variant) were found. It is known that during the agrobacterial transformation T-DNA was integrated in random regions of the plant genome (Mayerhofer 1991; Neve 1997), because the integration mechanism is not site-specific. Thus, in this case the multicopy of cT-DNA is not associated with successive transformation acts, as in *Nicotiana* species containing several cT-DNA, but with the duplication of cT-DNA together with the border regions of plant DNA after integration. Since cT-DNA in *Linaria vulgaris* was integrated into the transposon (Matveeva et al. 2012 and MH298319 sequence), a possible reason for cT-DNA duplications could be a temporal activation of this type of transposon which caused a transposition of its cT-DNA-containing copy. In our opinion the formation of extensively distant variants, can be explained by an unequal crossing-over which leads to the loss of individual elements (Fig. 3)

**Figure 3.**
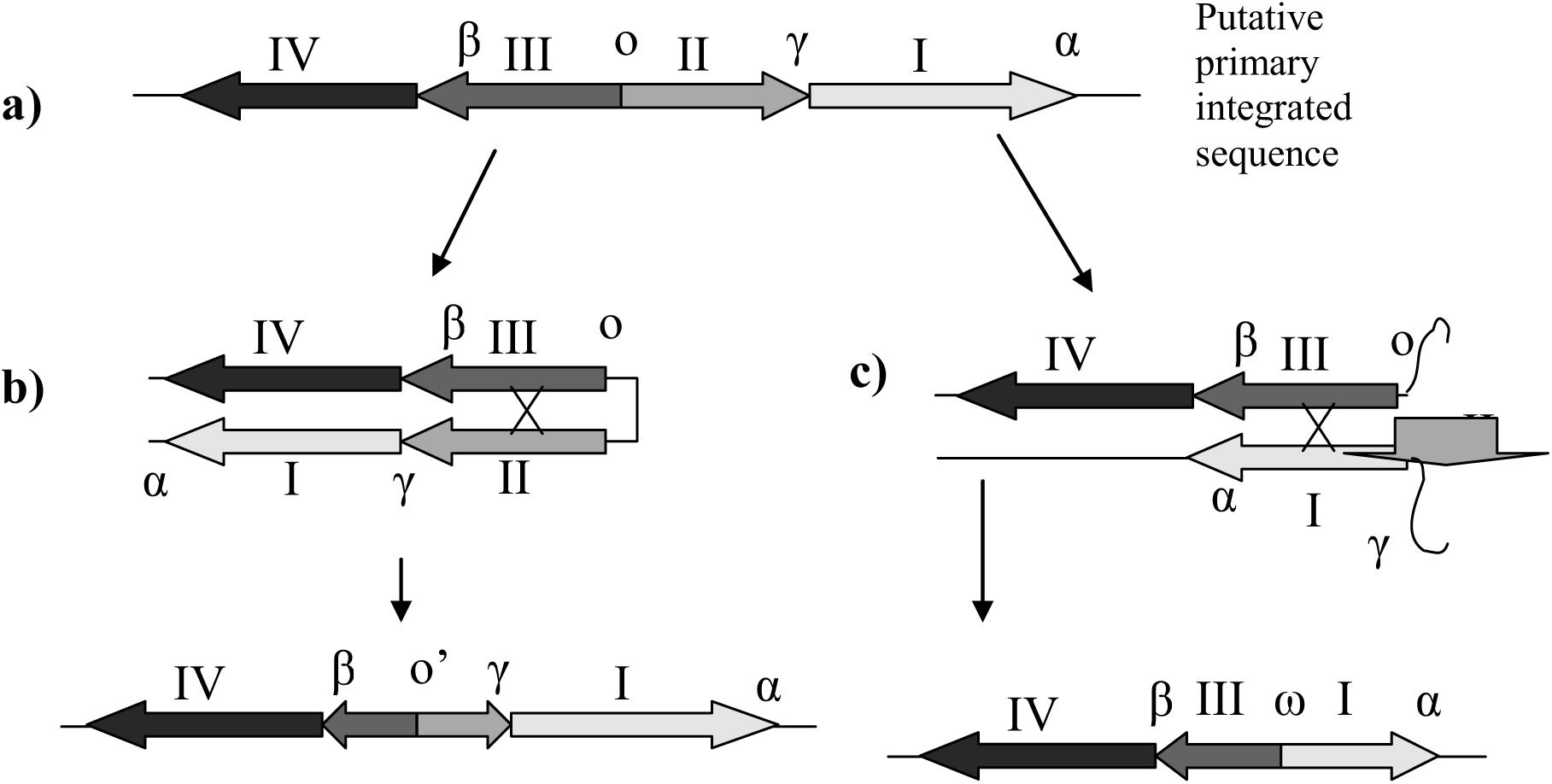
Possible unequal crossing-over can explain the occurrence of nested deletions in the region of o (b) and the formation of ω (c).

Potentially intact copies of the *orf13a* gene were detected in sequences 20.1, 9.13 and 206.2. The *orf13a* may encode the transcription factor (Hansen et al. 1994). Unfortunately, its function is not sufficiently studied.

Comparison of the sequences of different elements reveals, that a number of elements has the same large deletion of *rolC-mis*. This deletion exists in elements located in different positions, for example, I (20.4), II (20.5). Also we have found out, that this deletion is absent in the elements located in similar position (160.1, 18.2). The random occurrence of such deletions with accurately matching boundaries several times seems to be too great assumption. Explanation of the phenomenon due to illegitimate recombination with displacement requires double cross-over which is unlikely at such a short distance.

We assume that the cT-DNA elements undergo (or at least underwent earlier) he mechanism of homogenization (or “concerted evolution”). This process is similar to the mechanism, which aligns repeats of ribosomal RNA and other multicopy sequences (Pavelitz and Rusche 1995; Liao 1999) and maintains their high identity. Distribution of the variants with large deletions this way has also been described (Pavelitz and Rusche 1995).

Another evidence in favor of this hypothesis is the distribution of SNP-polymorphism in T-DNA elements, which cannot be explained by divergent evolution. Phylogenetic trees, constructed on the base of different SNP-sites, contradict each other (Fig. 4). The alignment looks as if elements sequences were copied by fragments of 25-50 bp in size. This picture cannot also be explained by PCR-artifacts, such as chain change, since the elongation time in amplification programs was enough (see Materials and methods). Also mosaic patterns are observed at very short intervals (25-50 bp). However if, we assume, that this is a chain change, then it would have to occur in 25-50 bp steps, which is impossible. Thus, it is impossible to give any alternative hypotheses explaining the observed phenomenon, except concerted evolution.

**Figure 4.**
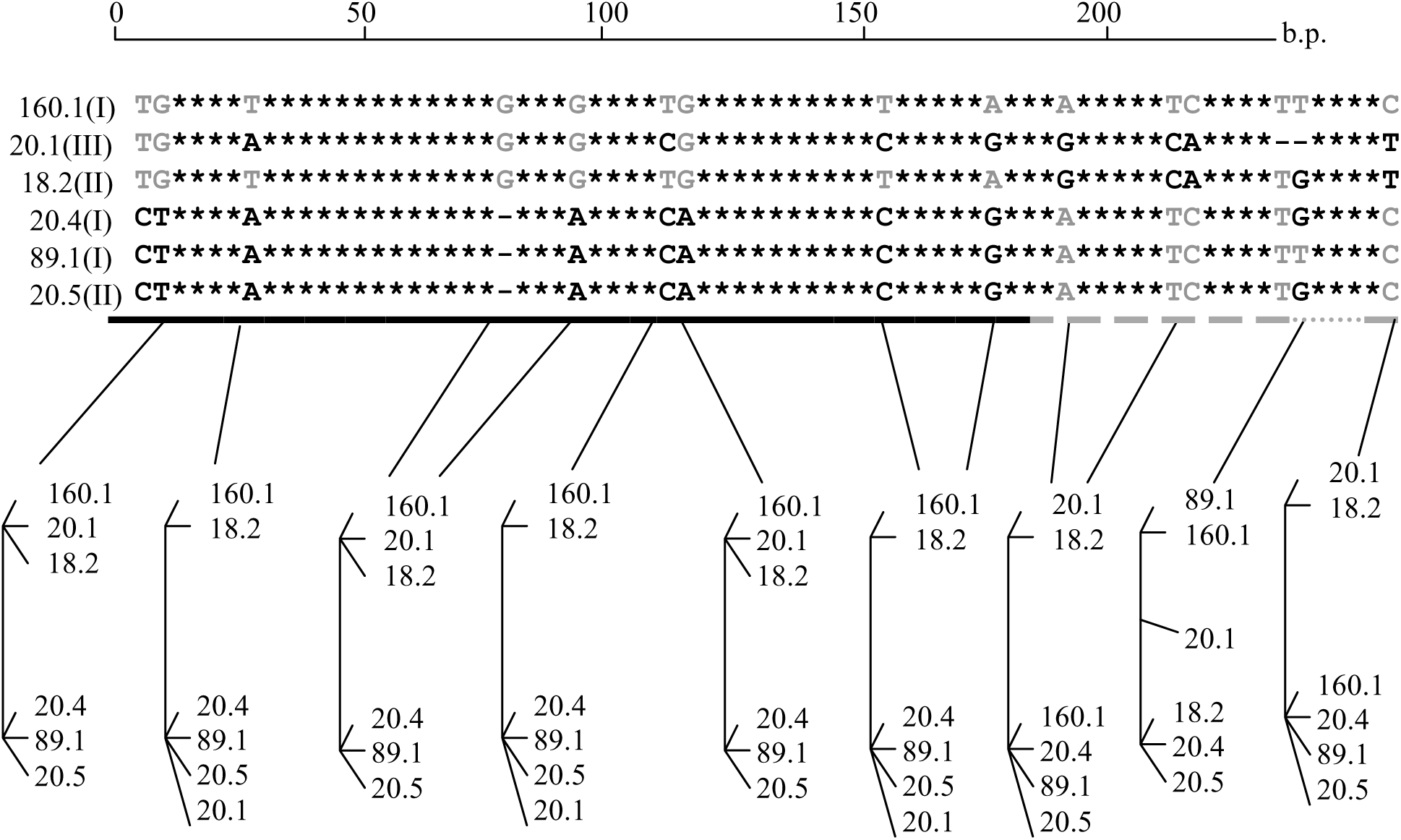
Alignment of *mis* gene fragment in various studied elements (only polymorphic sites are shown). Phylogenetic trees, constructed on the base of different polymorphic sites, contradict each other. Bold lines (solid and dotted) underline polymorphic positions, which give phylogenetic trees of different types.

It seems that the mechanism of concerted evolution works efficiently on high-copy and not very divergent sequences (Liao 1999). Typical examples of such sequences (rDNA, U2 RNA, histone genes) used as models of concerted evolution usually satisfy these requirements. Consequently, forthis reason the homogenization process is often observed at final stages, when the alignment has already been completed. In case of *Linaria vulgaris* cT-DNA, individual elements of cT-DNA have undergone less degree of homogenization and it is possible to observe an incomplete state of the process. Particularly, the discreteness of the process is visible.

According to our data, homogenization is progressing due to the replacement of short (25-50 bp) fragments of the sequence, but not whole repeat elements. *Linaria vulgaris* cT-DNA can become a valuable model object for studying the process of concerted evolution.

## Conclusions

According to the results, the following conclusions were made:

The structure of cT-DNA in *Linaria vulgaris* has been studied at a deeper level. According to new data, cT-DNA elements form inverted repeats (which is now shown for all cT-DNA-containing plants). The cT-DNA unit is organized as a complex inverted repeat of two simple tandem repeats. This block was duplicated at least twice after integration into the genome. Elements of cT-DNA are subject to homogenization process.

## Materials and methods

DNA was isolated from *L.vulgaris* leaves by CTAB-method (Doyle and Doyle 1987).

Long-range PCR (with amplicon size >2 kb) were made with a set of “D-polymerase” (Beagle Ltd., Russia) with a mixture of *Taq* and *Pfu* polymerases.

A set of the specific primers for junction sites was designed (see Fig. 1, Table 1) to exclude the possibility of their non-targeted annealing to other elements. Also, 10 forward (Set of universal forward primers for T-DNA elements, (see Table 1) and 26 reverse universal primers (Set of universal reverse primers for T-DNA elements, (see Table 1) were designed for the internal non-unique regions of the elements (Fig. 5). They were used to test specific primers, and to determine the approximate boundaries of the elements.

**Figure 5.**
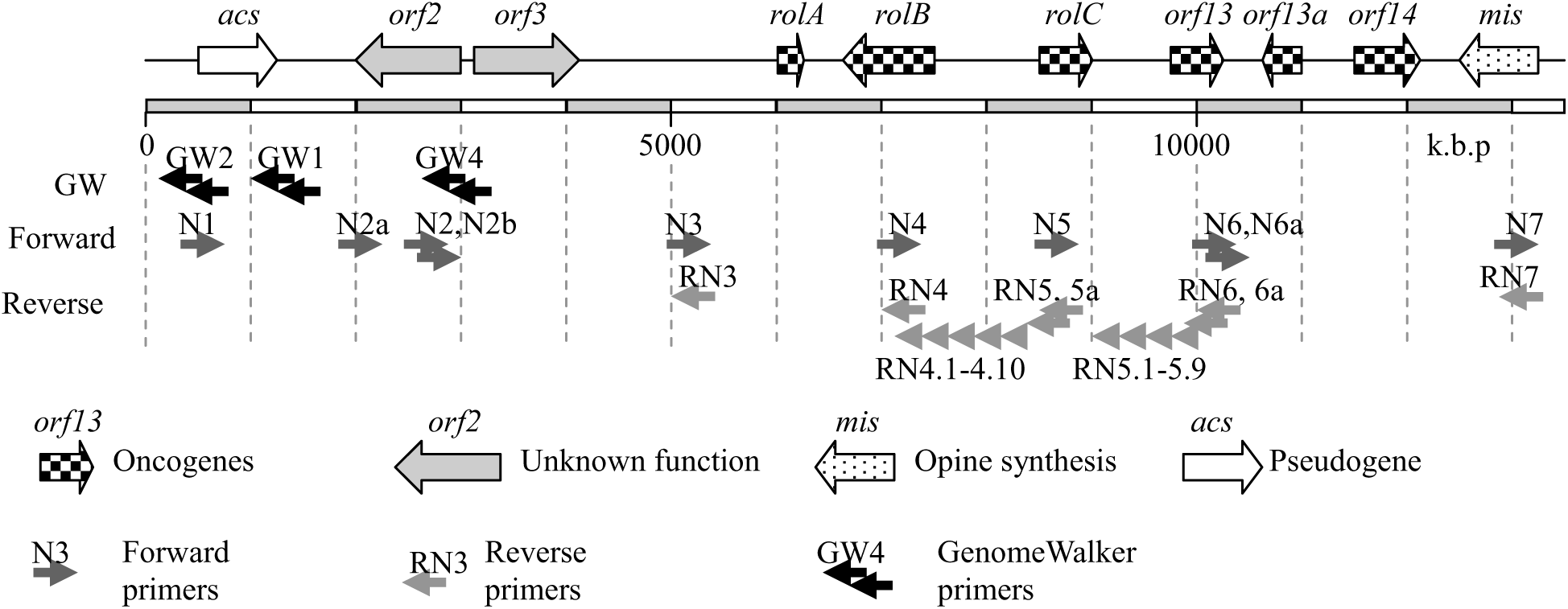
Positions of GemomeWalker primers and primers for non-unic regions of T-DNA element.

Reaction mixture consisted of 1x Buffer for D-polymerase, 0.3 mM dNTP, 0.5 μM forward primer, 0.5 μM reverse primer, 0.125 U / μl D-polymerase. The reaction was performed in a volume of 20 μl with 1 μl of plant DNA in PTC-100 thermocycler (MJ Research) with step-down protocol: 94 °C - 3 min, 7 cycles with high annealing / elongation temperature (94 °C - 5 s, 70 °C - 10 min), 38 main cycles (94 °C - 5 s, 65 °C - 10 min), and final elongation at 72 ° C for 7 minutes. PCR results were evaluated by electrophoresis in a 0.5% agarose gel on 1x SB buffer (Goedhart et al. 2005).

Sequencing was performed on a MegaBACE 1000 sequencer (Molecular Dynamics) using the manufacturer’s protocol.

Primer design was made for specific cT-DNA sites of junction between elements of repeats with each other or with the adjacent plant DNA (Fig. 1). Previously unknown junction sites were sequenced using the GenomeWalker approach (Siebert et al. 2005). The designed specific primers were checked for operability with a set of non-unique primers of the opposite direction, also the position of the adjacent element boundary was approximately determined.

The long-range PCR with various combinations of specific primers were conducted. Thus, the location of unique sites was studied (by the presence or absence of a PCR product), in the case of a positive result, a pure sequence of single elements was obtained after sequencing of a LongRange-PCR products.

To search for microsatellite markers *L. vulgaris* genomic DNA was sequenced with Ion Torrent System (TermoFisher Scientific, sequencing performed at Azco BioTech, USA). Obtained reads were used only for microsatellite motifs search, since a low coverage does not allow using this data for genome assembly or studying the structure of T-DNA. Digital PCR was performed by Azco BioTech (USA).

## Authors’ contributions

Conceived and designed the experiments: BD, IV OP.

Conducted the experiments: BD, IV OP.

Performed the analysis: BD, IV OP, DP.

Wrote the paper: IV, BD, OP.

All authors read and approved the final manuscript.

## Acknowledgements

The authors would like to thank Prof. Aleksandr V. Rodionov, Dr. Elena A. Andreeva, Dr. Yuliya V. Sopova, Dr. Mikhail Liskovykh, Dr. Maria S. Vishnevskaya, the Editor and Anonymous reviewers whose comments greatly improved this manuscript.

